# Five millennia of *Bartonella quintana* bacteremia

**DOI:** 10.1101/2020.03.13.990796

**Authors:** Ba-Hoang-Anh Mai, Rémi Barbieri, Thomas Chenal, Dominique Castex, Richard Jonvel, Davide Tanasi, Patrice Georges-Zimmerman, Olivier Dutour, David Perissinotto, Coralie Demangeot, Michel Drancourt, Gérard Aboudharam

## Abstract

*Bartonella quintana* caused trench fever in the framework of two World Wars and is now recognized as an agent of re-emerging infection. Many reports indicated the popularity of *B. quintana* exposure since the 90s. For evaluating its prevalence in ancient populations, we used real-time PCR to detect *B. quintana* DNA in 400 teeth collected from 145 individuals dating from the 1st to 19th centuries in nine archeological sites with the presence of negative controls. Fisher’s exact test was used to compare the prevalence of *B. quintana* detection in civil and military populations. *B. quintana DNA* was confirmed in a total of 28/145 (19.3%) individuals, comprising 78 citizens and 67 soldiers, 20.1% and 17.9% of which were positive for *B. quintana* bacteremia, respectively. This study collected previous studies on these ancient samples and showed that the presence of *B. quintana* infection followed the course of time in human history; a total of 14/15 sites from five European countries had a positive prevalence. The positive rate in soldiers was higher than those of citizens, with 20% and 18.8%, respectively, in the 18th - 19th centuries, but the difference in frequency was not significant. These results confirmed the role of dental pulp in diagnosing *B. quintana* bacteremia in ancient populations and showed the incidence of *B. quintana* in both citizens and soldiers. Many recent findings contributed to understanding the coevolution of the relationship between *B. quintana* and humans.

## Introduction

In June 1915, a British military doctor on the Western Front of the First World War reported the first case of recurrent fever in a soldier presenting headache, dizziness, and severe pain in the lower back and leg ^1^. Additional cases were reported among soldiers in trenches, after which it was named trench fever; causative agent *Rickettsia quintana* isolated from a patient in Mexico City by Vinson in 1961 was reclassified in genus *Rochalimaea* ^2,3^ and as *Bartonella quintana* in 1993 ^4^. The human body louse (*Pediculus humanus corporis)* is the transmission vector. *B. quintana* colonizes the gastrointestinal tract, is shed in louse feces, penetrates the human body through altered skin and enters the bloodstream ^5,6^. Clinical forms of the infection include trench fever, chronic bacteremia, endocarditis, bacillary angiomatosis, and lymphadenopathy ^5–7^. Since the 1990s, *B. quintana* has been recognized as a re-emerging agent in homeless populations in relation to unsanitary living conditions^8–11^.

The presence of trench fever was related to the military in Europe before the First World War through the finding of *B. quintana* DNA sequences in dental pulp and in body lice from soldiers of Napoleon in Vilnius (1812) ^12^ or in bone from soldiers in Kassel (1813-1814) ^13^. Indeed, dental pulp with high vascularity has been the definitive key in diagnosing bacteremia caused by microorganisms in paleomicrobiology ^14^. Molecular biology studies have confirmed *B. quintana* bacteremia occurring 200 to 4,000 years ago by the identification of *B. quintana* in dental pulp ^15^. There has been no comprehensive study of the prevalence of the pathogen in the past two millennia in Europe. To obtain such a figure, we reviewed all paleomicrobiological data regarding *B. quintana* and added the study of 400 additional specimens collected from 145 individuals over 20 centuries.

## Results

A total of 400 ancient teeth of 145 individuals collected in nine European sites with different times of burial were analyzed by qPCR. All negative controls were negative, and qPCR detected *B. quintana* DNA at all sites. The results indicated that a total of 78 citizens from six sites had 17.9% positivity, and 67 soldiers from three sites had 20.1% positivity. The positive detection of the Besançon site was the oldest, with data from the 1st-4th centuries (Table S1). Seven of 17 archeological sites had data from the 18th-19th centuries (Tables S1 and S2), where the prevalence of *B. quintana* DNA was 3/16 (18.8%) among citizens and 24/120 (20%) among soldiers; the difference in the prevalence was not significant (Table S3).

## Discussion

The results reported here were authentic. Indeed, we chose ancient teeth with a closed apex that were free of traumatic lesions, which helped avoid any risk of external contamination, and a positive control was not used in PCR experiments because it could be a source of contamination; the negative controls remained negative ^16^. For screening a total of 400 teeth, we used real-time PCR as previously described to identify *B. quintana* ^16–18^, and in this work, each experimental step was performed in different rooms of the new building where these gene sequences had not been previously used. The number of teeth per individual was variable in each burial site because of availability, and we used as many teeth per individual as possible to increase the chance of detection. Indeed, the first report of using dental pulp for ancient septicemia diagnosis indicated that the number of teeth per individual infected with *Y. pestis* was 1/4 teeth, 1/2 teeth, and 3/3 teeth ^14^; furthermore, two teeth from one individual reported that the first infected *Y. pestis* and the second had co-infection of *Y. pestis* and *B. quintana* ^16^.

The history of trench fever was marked in 1915 by reporting new clinical features from soldiers in trench warfare ^2,3^. In 2005, authentic evidence showed that *B. quintana* caused bacteremia in humans for more than 4000 years by DNA analysis in dental pulp collected from Peyraoutes-France ^19^.

Afterward, this pathogen was also identified in this specimen from different burial sites, including Venice (15th-16th) ^18^, Douai (18th) ^17^, Vilnius (1812) ^12^ and Bondy (11th-15th) ^16^, where an individual had coinfection with *Y. pestis*. In addition to dental pulp, *B. quintana* was detected in ancient bone (1813-1814) ^13^ and coprolites (14th) ^20^. Bacterial confirmation in ancient pulp indicates that individuals had bacteremia before death, but it might not be the cause of death, proving that no mortality has been recognized as being caused by trench fever ^5^. According to a survey of 930 homeless people in Marseille, 5.3% were blood culture positive for *B. quintana* ^8^, and asymptomatic chronic bacteremia could be maintained for 78 weeks ^21^. In 2004, *B. quintana* was found in the dental pulp of a homeless patient having bacteremia the previous six months ^22^.

The finding of *B. quintana* in dental pulp comes mostly from our laboratory, with nearly half of burial sites located in the 18th-19th centuries; 7/14 sites were in France, 4/14 were in Italy, and the rest came from Ukraine, Russia, and Lithuania. This can be explained by the availability of samples in these periods, and our research team located in France had many favorable conditions to receive these teeth from archeological centers. The wide distribution of geography and time suggested that this bacterial infection was common in historic European populations. In the medical literature, the “five-day fever” and the Moldavia fever in the 19th century had clinical signs similar to trench fever but lack of laboratory techniques for confirmation ^23^.

Millions of soldiers were contaminated worldwide during two world wars but after the end of each war, the incidence dropped dramatically but still sporadically persisted in some countries ^2^. It was noted that approximately 20% of Napoleon’s soldiers in Vilnius, Lithuania, were exposed to body lice containing *B. quintana* in 1812 ^12^. An investigation of ancient bones of 18 soldiers from a historical mass grave inhumed in winter 1813/14 in Kassel, Germany, revealed that 16.7% of soldiers had the infection ^13^. In our study, soldiers from Kaliningrad site had only 6.7% positive results, in contrast with higher positive rate of Dax or Sevastopol, this can be explained by the use of only one tooth per individual for Kaliningrad samples. Catacombs of St. Lucia had the similarity of almost using one tooth with the lowest positive proportion (3.6%), this site represented as the ancient Christian monument of Late Roman period ^24^.During war, soldiers were generally crowded closely together in unsanitary conditions for prolonged periods of time; moreover, despite knowing the role that lice played in transmission and their presence in clothes, there were no effective methods of disinfection available ^2,3^.

To have a homogeneous time, we chose the 18th-19th centuries with a greater number of both populations. The incidence in citizens and soldiers was 18.8% and 20%, respectively, and there was no significant difference. This suggests that this bacterial infection is relatively common throughout human society, or moreover, the sample size of citizens was small (18 citizens), which could affect the comparison. The epidemiology of trench fever involves interhuman transmission via vectors, such as body louse infection, other transmission that may contact reservoirs, bites of insects ^25^, bites of cats ^26^ or others that have yet to be defined and need to be further elucidated. Humans are not the only reservoirs of *B. quintana*; it is also found in cat fleas ^27^ and monkey fleas ^28^, and some studies have indicated that other animals as known reservoirs via detection of *B. quintana* in domestic cats ^29^, dogs ^30^, rhesus macaques ^31,32^, and Japanese macaques ^33^. Following an experimental study on cat fleas, this pathogen was absorbed via the gastrointestinal tract and released its into feces ^34^. Another study showed that the *Pedicinus obtusus* louse was postulated as an efficient vector of transmission between rhesus macaques ^32^. Homelessness was defined as a high-risk factor for *B. quintana* infection as a consequence of inadequate hygiene ^8,10,11,21^; however, some subpopulations have relatively high exposure rates, such as blood donors, between 27% and 51%, and ^9,35,36^ healthy people, between 11.2% and 25% ^36–39^. There is no evidence to assume that these people had hygienic living conditions, such as homelessness, or contacted body lice infection, thus suggesting the existence of some underestimated factors that cause a bacterial infection. With the new transmission findings, transmitted vectors, reservoirs and widely infected populations contribute to the understanding of the infection from the past to the present.

## Methods

### Ancient human samples

Ancient teeth were collected from human remains in nine European burial sites by archeologists and transported to our laboratory (Supplementay 1). Following our current protocol for teeth choice and their manipulation ^18^, teeth were washed with sterile water and gradually dried, and the dental pulp was extracted using rotating disc instruments as previously described ^40^. Total DNA was extracted by using the phenol-chloroform protocol ^41^.

### Molecular detection

Dental pulp was tested for *B. quintana* DNA using quantitative real-time PCR (qPCR) targeting the ITS gene using the following primers and probes: probe/6FAM-GCG CGC GCT TGA TAA GCG TG-TAMRA, forward/5’-GAT GCC GGG GAA GGT TT TC-3’, reverse/5’*-*GCC TGG GAG GAC TTG AAC CT-3’ ^16–18^. qPCR amplification was performed using the LightCycler® 480 Probes Master Kit according to the manufacturer’s recommendations (Roche Diagnostics, Meylan, France). Each well contained 10 µL of mix, 3 µL of sterile water, 0.5 µL of probe (20 µM), 0.5 µL of each primer (50 µM), 0.5 µL UDG and 5 µL of extracted DNA. Amplification consisted of a 2-min incubation step at 50°C and an initial 5-min denaturation at 95°C, followed by 40 cycles of denaturation at 95°C for 5 seconds and hybridization at 60°C for 30 seconds. Two negative controls (control of DNA extraction + mix and sterile water + mix) were placed every 5 samples in each plate. A sample was considered positive when the qPCR result was positive with a cycle number (Ct) lower than 40.

### Fisher’s exact test

Fisher’s exact test was used to compare the prevalence of detected *B. quintana* in the civil population and soldier population, and a p-value <0.05 was considered statistically significant.

### Conclusions

Dental pulp is a favorable sample for investigating *B. quintana* bacteremia in ancient populations, and it is relatively easy to collect and is well protected inside teeth. Previous studies collected in this study showed the presence of *B. quintana* infection 4,000 years ago and from the 1st to 19th centuries. Soldiers are considered the main target of this bacterial infection in human history, but we observed in this study no significant difference between citizens and soldiers during the 18th-19th centuries.

## Supporting information

Supplementary Informations

## Acknowledgments

This work was supported by the French Government under the Investissements d’avenir (Investments for the Future) program managed by the Agence Nationale de la Recherche (ANR, fr: National Agency for Research), [reference: Méditerranée Infection 10-IAHU-03]. This work was supported by Région Le Sud (Provence Alpes Côte d’Azur) and European funding [FEDER BIOTK].

- Thanks to Michael Decker (University of South Florida, Department of History, Tampa, Florida,USA) for his information on the samples.

- Thanks to the Pontifical Commission for Sacred Archaeology - Inspectorate for the Catacombs of Eastern Sicily for the permit to work other samples from the Catacombs of St. Lucy art Siracusa - Italy

## Author contribution statement

B-H-A.M. performed experiments, interpreted the data and drafted the manuscript.

R.B. received, managed samples, performed some experiments, drafted the manuscript.

T.C. collected samples, described archaeological site, drafted the manuscript.

D.C. collected samples, described archaeological site, drafted the manuscript.

R.J. collected samples, described archaeological site, drafted the manuscript.

D.T. collected samples, described archaeological site, drafted the manuscript.

P.G-Z. collected samples, described archaeological site, drafted the manuscript.

O.D. collected samples, described archaeological site, drafted the manuscript.

D.P. collected samples, described archaeological site, drafted the manuscript

C.D. collected samples, described archaeological site, drafted the manuscript.

M.D. conceived the experiment, interpreted the data and drafted the manuscript.

G.A. conceived the experiment, interpreted the data and drafted the manuscript.

## Competing interests

We declare no competing interests.

## LEGENDS

**Table 1:** Results of molecular biology to detect *B. quintana* in ancient dental pulp.

| Sites | Datation | Positive teeth/total | Positive people/total | Number of teeth/person | Positive percentage | Population |
| --- | --- | --- | --- | --- | --- | --- |
| Besançon - France | 1st - 4th | 7/29 | 3/5 | 4-7 | 17.9% (14/78) | Citizens |
| Catacombs of St. Lucia - Italia | 3th - 6th | 1/29 | 1/28 | 1-2 |  |  |
| San Basilio - Italia | 4th - 6th | 1/8 | 1/6 | 1-2 |  |  |
| Dueville - Italia | 6th - 7th | 2/15 | 2/15 | 1 |  |  |
| Remiremont - France | 5th - 10th | 13/45 | 4/8 | 3-8 |  |  |
| Amiens - France | 18th - 19th | 4/55 | 3/16 | 3-4 |  |  |
| Dax - France | 1792-1833 | 11/110 | 4/9 | 9-14 | 20.1% (14/67) | Soldiers |
| Kaliningrad - Russia | 1812 | 2/30 | 2/30 | 1 |  |  |
| Sevastopol- Ukraine | 1853-1856 | 14/79 | 8/28 | 1-5 |  |  |

**Table 2:** *B. quintana* detection in ancient specimens from previous studies.

| Sites | Datation | Specimens | Methods | Positive number<br>of specimens/total | Positive number<br>of people/total | Population | Ref. |
| --- | --- | --- | --- | --- | --- | --- | --- |
| **Peyraoutes - France** | 2000BC | Dental pulp | Suicide PCR ( groEL, hbpE genes ) | 1/6 | 1/3 | - | (Drancourt et al.,<br>2005) |
| **Roaix - France** | 2200BC - 2100BC |  |  | 0 | 0/3 |  |  |
| **Bondy - France** | 11th - 15th |  | Real time PCR (ITS gene ) | 3/14 | 3/5 | Citizens | (Tran et al.,<br>2011a) |
| **Venice - Italia** | 15th - 16th |  |  | 5/173 | - | Citizens | (Tran et al.,<br>2011b) |
| **Douai - France** | 18th |  |  | 1/40 | - | Soldiers | (Nguyen-Hieu et<br>al., 2010) |
| **Vilnius - Lithuania** | 1812 |  | Suicide PCR (hbpE, htrA genes) | 7/72 | 7/35 | Soldiers | (Raoult et al.,<br>2006) |
| **Kassel -Germany** | 1813-1814 | Bone | PCR standard ( hbpE gene) | 3/18 | 3/18 | Soldiers | (Grumbkow et al.,<br>2011) |
| **Namur - Belgium** | 14th | Coprolite | 16S rRNA metagenomics |  |  |  | (Appelt et al.,<br>2014) |

(-): Not mentioned

**Table 3:** Comparison of infected populations in the18th - 19th centuries.

|  | **Total** | **Positive number** | **Percentage** | **p** |
| --- | --- | --- | --- | --- |
| **Citizens** | 16 | 3 | 18.8 % | > 0.05 |
| **Soldiers** | 120 | 24 | 20 % |  |

