## Supplementary Informations for "Five millennia of *Bartonella quintana* bacteremia"

**Nine** **European archeological sites.**

**1. Remiremont - France**

Saint Mont was the first rural monastery in Austrasia. Between 2014 and 2017, a major funeral building was discovered. Many formal burials were discovered in a building of a real Merovingian funerary complex installed at the top of a mountain. A total of 650 kg of bones were found in a tertiary pit, which is located on the slopes adjacent to the site. The complex excavation of these remains from an ossuary suggested that a population linked to the monastery, dated from 7th to 8th century, was buried there, but everything revealed that medieval populations have benefited from what was certainly the funeral church of the monastery.

**2. Besançon - France**

During the construction of the Viotte railway station-Besançcon in Eastern France in the 19th century, a vast sepulchral space was discovered with many objects and human remains dating to the Gallo-Roman era. In 2012, the surface of seven trenches was excavated for further investigation after the excavation operation of the whole space was realized. The results showed forty-eight funerary structures with 43 skeletons that were more or less well preserved and had funerary structures, such as ceramics, glass, fauna, and shoes. By many methods of archeological identification, all the funerary structures were dated to the 2nd to 4th century.

**3. Dax - France**

Upstream of a real estate project on the right-of-way of Lycée Saint-Joseph in Dax (Landes), on the site of the former Capuchin convent founded in 1614 and converted into military barracks in 1823, preventive excavations were carried out by Hades (RO, N. Sauvaitre, Anthropologists: D. Peressinotto and C. Demangeot). Multiple burials have been discovered within the burial space. They consisted of several pits arranged in a row with an increasing number of individuals going from east to west. The positions of the subjects and the taphonomy evoked bodies naked thrown in the pits, without archeological artifacts. The sample consisted of exclusively male subjects, the majority of whom were under 30, with some slightly older individuals, but not older than 40 years of age. The assumption is that they are soldiers in the field, victims of a health crisis, in particular, because of the total absence of war wounds and the management of the bodies.

**4. Amiens - France**

The archeological site of the rue de la Résistance in Amiens unearthed the remains of the Hotel-Dieu of the city, built in the 14th century and active until the 18th century. This former hospital was, in medieval times, on an island surrounded by canals. This search also revealed three pits located in the cemetery area of this building: multiple burials containing several dozen individuals buried in the shroud and deposited in a very short time. Radiocarbon dating from buried individuals indicated that these pits dated back to the 18th century.

**5. Dueville - Italy**

Dueville is located approximately 12 kilometers north of Vicenza in the Belvedere district. In 1911, during roadwork for a new road, the Dueville-Montecchio-Precalcino-Langobard tombs were discovered. Dueville was excavated again in multiple campaigns beginning in 1993 through 2009 by the Veneto Archeological Superintendence, which uncovered more than 500 burials, making it one of the largest Langobard necropolises in Italy ^1,2^. Remains from these excavations were radiocarbon dated the 7th-9th centuries AD; however, the artifacts associated with many of the burials indicated the early migrating Langobard culture during the late 6th to 7th century AD ^2^. Carrara reported on the burial context for excavations from 2000 to 2009 in which 217 burials were excavated. Primary burial pits were oriented west-east with heads to the west. Individuals were supine, with variations in the upper limbs including along the sides, one on the side and the other on the chest or pelvis, and arms crossed on the chest or pelvis ^2^. Most of the burials were directly in the ground, while 7.8% were buried in coffins, and 19.8% showed evidence of the use of shrouds. Skeletal remains from the excavated site of Dueville (2002, 2005, 2008) were housed at the Palladio Museum in Vicenza and sampled for the present research study.

**6. San Basilio (Rovigo) - Italy**

Archeological excavations in San Basilio uncovered a large Roman building comprising an early Christian church with an associated cemetery, well and additional structures from the Early Medieval Period. The necropolis was used from the 4th to 6th century AD and correlated with the different building stages of the church. It expanded into previously abandoned structures. Thirty-four tombs were identified, including children, and their location near the church suggested the importance of the religious center at the time. The majority of the burials were oriented west-east; however, a few were buried north-south potentially to conserve space in the necropolis or establish a higher social position as they were located against the church. Many of the burials were catacomb style, with brick roofs.

**7. Catacombs of St. Lucia - Italy**

The Catacombs of St. Lucia represent one of the oldest and most important monuments related to the Christian communities of Syracuse and Sicily in the Late Roman period ^3^.Beneath the homonymous square, there was a large underground cemetery that developed from the 3rd to the 5th century AD, incorporating previous structures used for funerary, cultural and industrial purposes that were transformed into monumental burial chambers. The presence of the tomb of St. Lucia guaranteed the popularity of the complex even after the end of its use as a cemetery in the 6th century AD. In fact, in at least two regions of the catacombs, new oratories were built, probably relating to the activity of nearby monastic groups. Sector F of Region C, also known as the “second pagan shrine”, belonged to the pre-existing structures of the late Hellenistic and Roman periods, later incorporated in the cemetery. A hypogeal shrine originally designed around a pillar carved in the rock and decorated with niches was drastically transformed through very complex phases of use, which radically altered its original appearance. Region C of the catacombs, definitively dated to post-Constantinian times (4th century AD) and used for funerary purposes until the end of the 5th century AD, offered significant evidence for the Medieval phase of use. After the cemetery fell into disuse for burial, the catacombs underwent a series of architectural and monumental transformations that turned it into a true place of martyrial worship: in particular, the so-called Oratory C, Crypt VI comprised a group of cubicula set on different levels, accessible from a single monumental entrance with a sequence of three framed arches. In a further phase, a lower level connected to the rest of the complex by a staircase was excavated. Portions of frescoes related to the last phase of use are still visible on the walls. In one room, there were some massive sarcophagi carved into the bedrock, indicating the high rank of the commissioner of this complex. In the framework of the research project on dietary habits and health style of the Christian community buried in the Roman Catacombs of St. Lucy (P.I. Davide Tanasi, University of South Florida; Co-P.I. Gioacchina Tiziana Ricciardi, Pontificia Commissione di Archeologia Sacra, Ispettorato per le catacombe della Sicilia Orientale) on October 2017, 64 samples (38 teeth and 26 bone fragments) were collected from a large number of individuals in burials excavated between 2011 and 2015 in Sector F, Oratory C, and Crypt VI ^4^.

**8. Sevastopol - Ukraine**

At a construction site located at "33, street of Truth" (Севастополь, Ул Правды 33), in a suburb of Sevastopol, near Kamiesch Bay, individual burials of soldiers were found. These soldiers were French given the location where the French army stationed for the military seat of Sevastopol in 1853, and some evidence was found (buttons of jackets of the 11th and 39th regimens of infantry of line). An anthropological mission in September-October 2013 characterized the bones. This anthropological mission identified 99 individuals among whom there were only men, more than half of whom were under 40 years of age.

**9*.* Kaliningrad - Russia**

In July and August 2006, archeological excavations were carried out in downtown Kaliningrad. These excavations permitted the identiﬁcation of a part of the fortiﬁcations in the old town of Königsberg (the capital of Eastern Prussia) and the discovery of a military mass grave dated from late 1812 to early 1813, based on the buttons and fragments of uniforms of the Napoleon Grande Armée associated with the human remains. The mass grave was organized in two parallel rows, the skeletal assemblage was from 800 to 1,000 individuals, the sex ratio appears to be strongly male-biased, only 5% of sex-determined skeletons were female, and more than 80% of this sample belonged to the 15-25 year age category. Observation of traumatic lesions, including evidence of surgery, support the hypothesis that this mass grave contained soldiers bodies who died in military hospitals of city ^5^.

5. Dutour, O. & Buzhilova, A. Palaeopathological Study of Napoleonic Mass Graves Discovered in Russia. The Routledge Handbook of the Bioarchaeology of Human Conflict, Chapter 26. London: Taylor and Francis, p. 511-524.
